# Cell cycle dynamics of redox state and lipid metabolism in *S. cerevisiae*, *S. pombe* and murine leukaemia cells

**DOI:** 10.64898/2026.01.22.701002

**Authors:** Hanna M. Terpstra, Julius A. Fülleborn, Julia Kamenz, Matthias Heinemann

## Abstract

Coordination of metabolism, cell growth and cell division is essential to life. Recent single-cell measurements in *S. cerevisiae* have shown that metabolic processes and the cellular redox state are dynamic along the cell cycle. However, it is unknown whether similar metabolic oscillations also occur in other organisms. Until now, the dynamics of metabolism in other eukaryotes have predominantly been studied in cell cycle synchronised populations. Since cell cycle synchronisation methods can perturb metabolism, they may also introduce artefacts in the recorded dynamics. Here, we performed time-lapse microscopy analyses of exponentially growing single cells of the budding yeast *S. cerevisiae*, the fission yeast *S. pombe* and murine leukaemia L1210 cells. Measuring the NAD(P)H autofluorescence and the cell surface area growth rate in unsynchronised cells, we discovered oscillations along the cell cycle of the cellular redox state and lipid metabolism, respectively. Thus, our work shows that metabolism is dynamic along the cell cycle of these three evolutionarily distant eukaryotic organisms. This finding suggests that such metabolic oscillations could be a conserved characteristic among eukaryotes.

## Introduction

A growing body of research shows that *S. cerevisiae* metabolism is dynamic during the cell growth and division cycle. Metabolic dynamics related to the cell cycle were first observed in glucose-limited continuous cultures where cells can synchronise their cell cycles (Borzani et al., 1977; Finn & Wilson, 1954; Tu et al., 2005; von Meyenburg, 1973). These synchronised chemostat cultures revealed alternating periods of high and low oxygen consumption (Tu et al., 2005). The period of high oxygen consumption is coupled to cell cycle START (Burnetti et al., 2016; Shi & Tu, 2013; von Meyenburg, 1969). Changes in the oxygen consumption rate also are concomitant with periodic changes in gene expression (Klevecz et al., 2004; Tu et al., 2005) and metabolite levels (Tu et al., 2007). More recently, multi-omics studies using cell cycle-synchronised cultures have shown that the proteome and metabolome also are dynamic along the cell cycle (Blank et al., 2020; Campbell et al., 2020).

Moreover, metabolic oscillations along the cell cycle have been observed in single budding yeast cells. It was found that the protein synthesis rate peaks twice per cell cycle (Takhaveev et al., 2023), with its first peak shortly before START driving commitment to the cell division cycle (Litsios et al., 2019). Lipid biosynthesis peaks once per cell cycle, midway through S/G2/M (Takhaveev et al., 2023). The glycolytic flux is also dynamic, peaking around cytokinesis (Monteiro et al., 2019). Besides individual metabolic processes, the redox state of the cell, as measured through autofluorescence of NAD(P)H (Papagiannakis et al., 2017) or flavin autofluorescence (Baumgartner et al., 2018), was also found to oscillate on the single-cell level. It remains unclear whether similar cell cycle dynamics of metabolic processes exist in other eukaryotic organisms.

The available knowledge on the metabolic dynamics in eukaryotes other than *S. cerevisiae* is limited and was predominantly gathered on the population level. In elutriated cultures of the fission yeast *S. pombe*, a correlation between the periodic expression of metabolic genes and environmental stress genes was found, comparable to results in *S. cerevisiae*. This similarity indicates that *S. pombe* could go through metabolic cycles that are comparable to those in budding yeast (Slavov et al., 2012). Moreover, the protein synthesis rate in elutriated cultures of *S. pombe* has been shown to be dynamic (Creanor & Mitchison, 1982), as has the ATPase activity in cultures synchronised by continuous-flow size-selection (Edwards & Lloyd, 1977). In cultures synchronised by sedimentation-velocity size-selection, the dynamic abundance of tricarboxylic acid cycle enzymes was quantified (Poole & Lloyd, 1973) and two peaks of oxygen consumption per cell cycle were found (Poole et al., 1973). In addition, in cultures that were synchronised with 2’-deoxyadenosine, the cellular redox state was found to be dynamic (Bashford et al., 1980). While these findings suggest the existence of metabolic dynamics along the cell cycle in *S. pombe*, it is important to note that the applied synchronisation methods could have artificially induced these dynamics. For instance, sedimentation-velocity size-selection is a time-consuming procedure during which cells are subjected to suboptimal temperatures and anaerobic conditions, and chemically induced cell cycle synchronicity can interfere with metabolism and cause metabolic stress (Lloyd et al., 1975). Therefore, the observed metabolic dynamics could merely reflect a stress response to the cell cycle synchronisation methods. To the best of our knowledge, there are currently no single-cell data on the metabolic dynamics during the cell cycle of *S. pombe*.

In mammalian cells, our knowledge of the dynamics of metabolic processes along the cell cycle is similarly limited. Studies have shown that fluxes through central metabolic pathways, specifically the tricarboxylic acid cycle and glycolysis, measured on the population level in chemically synchronised cells, change along the cell cycle (Ahn et al., 2017; Liu et al., 2020). The ATP synthesis rate, as derived from the mitochondrial membrane potential, measured with a fluorescent probe in a single-cell buoyant mass sensor with a microscope mounted on top, also oscillates over the cell cycle (J. H. Kang et al., 2020). Furthermore, the NAD^+^/NADH ratio was found to be dynamic along the cell cycle in elutriated (Yu et al., 2009) as well as chemically synchronised (Ahn et al., 2017) cells. Yet, our understanding of metabolic dynamics during the mammalian cell cycle remains limited.

In this study, we asked whether metabolic oscillations on the single-cell level in unsynchronised, exponentially growing cells similar to those observed in *S. cerevisiae* also occur in *S. pombe* and murine leukaemia (L1210) cells. To this end, we used two label-free metabolic measurements, namely the NAD(P)H autofluorescence, a proxy for the cellular redox state as established in budding yeast (Papagiannakis et al., 2017), and the dynamic cell surface area growth rate, a proxy for lipid biosynthesis dynamics. We found that the NAD(P)H autofluorescence and the cell surface area growth rate are both dynamic along the cell division cycle in all three organisms. We also discovered that in *S. cerevisiae* and *S. pombe*, but not in L1210 cells, the derivative of the NAD(P)H autofluorescence signal anticorrelates with the cell surface area growth rate. This suggests that in these two organisms, lipid biosynthesis may have considerable effects on NAD(P)H levels. With this work, we revealed metabolic dynamics on the single-cell level in three eukaryotes, suggesting that metabolic dynamics during the cell cycle might be a shared characteristic of eukaryotic organisms.

## Results

### Analysis of NAD(P)H autofluorescence and cell surface area growth rate along the cell cycle

To elucidate dynamics of NAD(P)H autofluorescence and cell surface area growth rate over the cell cycle, we developed an analysis pipeline that starts with the collection of single-cell data using time-lapse microscopy. To obtain cell masks, we segmented bright-field images with YeaZ (Dietler et al., 2020), an algorithm originally developed to segment images of *S. cerevisiae*, which can also successfully segment images of *S. pombe* and L1210 **(Figure S1)**. These cell masks were then applied to the fluorescence images in order to determine the average cellular fluorescence intensity. Moreover, we calculated the cell surface area based on the cell masks’ dimensions and subsequently determined the cell surface area growth rate by dividing the difference in cell surface area between two subsequent time points by the imaging time interval **(Figure 1A)**.

**Figure 1.**
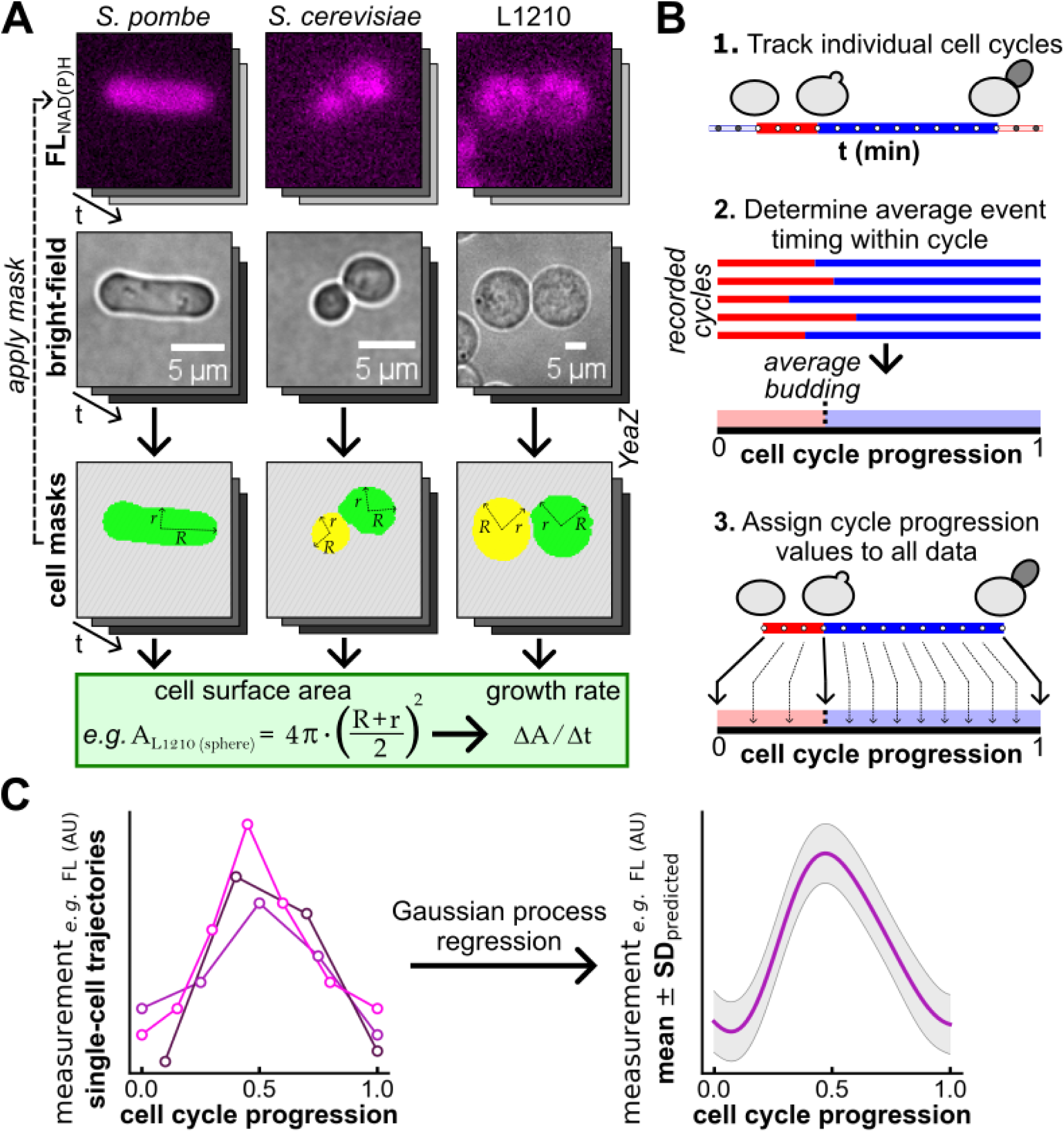
Analysis of NAD(P)H autofluorescence and cell surface area growth rate along the cell cycle. **(A)** Time-lapse microscopy of single cells in bright-field and NAD(P)H autofluorescence channels. Cell masks, generated from bright-field images with YeaZ, define regions of interest and also allow the calculation of the cell surface area, as well as its growth rate; **(B)** Single-cell trajectories are segmented into cell cycles based on events visible in the bright-field images, *e.g.* cytokinesis and budding in *S. cerevisiae*. Trajectories are mapped onto a common cell cycle progression coordinate, ranging from 0 to 1, by aligning the occurrence of these cell cycle events across cell cycles; **(C)** Gaussian process regression is applied to the aligned single-cell cycle data to predict the mean signal and associated standard deviation along the cell cycle. This yields the cell cycle dynamics of the NAD(P)H autofluorescence and the cell surface area growth rate in the ‘average cell’.

We monitored cell cycle progression of individual cells based on cell cycle events that are visible in the bright-field images, *i.e.* cytokinesis and budding in *S. cerevisiae*, septation, cell birth, which we defined as the moment of separation between the two daughter cells, and the appearance of the forming septum in *S. pombe*, and cytokinesis in L1210 cells. We defined the end of one cell cycle and the beginning of the next one as the moment of cytokinesis in *S. cerevisiae* and L1210, and septation in *S. pombe* **(Figure 1B; step 1)**.

To enable comparison between cell cycles of different durations, we projected the single-cell trajectories for NAD(P)H autofluorescence and the cell surface area growth rate onto a common cell cycle progression coordinate. On this coordinate, every cell cycle begins at time value 0 and ends at 1, and the occurrence of cell cycle events within the cell cycle, such as budding in *S. cerevisiae*, is aligned. To determine the cycle progression value for these events, we first calculated the fraction of each individual cycle that had elapsed when the event was detected. Then, we used the data from all recorded cycles to calculate the average cycle progression value at which this event occurred. This value was then assigned to all data points at which the cell cycle event was detected **(Figure 1B; step 2)**. After obtaining the cell cycle progression values for the data points at which a cell cycle event was recorded, we determined the cell cycle progression values for all data points recorded between cell cycle events. To this end, we evenly distributed these data points over the interval bounded by the two cell cycle events **(Figure 1B; step 3)**. Finally, after all single-cell trajectories had been projected onto the common cell cycle progression coordinate, we applied Gaussian process regression to the cell cycle-aligned data to predict the mean value and its associated standard deviation along the cell cycle **(Figure 1C)**. This analysis pipeline allowed us to determine the cell cycle dynamics of both the NAD(P)H autofluorescence and the cell surface area growth rate in the ‘average cell’.

### NAD(P)H autofluorescence and cell surface area growth rate oscillate over the *S. cerevisiae* cell cycle

First, we set out to confirm the validity of our data analysis pipeline by applying it to *S. cerevisiae*, whose cell cycle oscillations of NAD(P)H autofluorescence and cell surface area growth rate have been reported previously (Papagiannakis et al., 2017; Takhaveev et al., 2023). Applying the same acquisition and analysis pipeline to *S. cerevisiae*, *S. pombe* and L1210 allowed us to validate the previous findings, and also enabled a direct comparison between the different organisms with regard to the cell cycle dynamics of the NAD(P)H autofluorescence and the cell surface area growth rate later in this work.

In our time-lapse microscopy data of *S. cerevisiae*, we monitored cell cycle progression by detecting cytokinesis and budding in the bright-field images. Cytokinesis demarcates the end of one cell cycle and the beginning of the next, while budding denotes the G1/S transition **(Figure 2A)**. Following a single *S. cerevisiae* cell over three consecutive cell cycles, we observed oscillations of both the NAD(P)H autofluorescence and the cell surface area growth rate: NAD(P)H autofluorescence peaks shortly after budding and shows a trough before cytokinesis **(Figure 2B)**. In contrast, the surface area growth rate is lowest around budding and peaks midway through the S/G2/M phase **(Figure 2C)**. These dynamics are also visible in the data aligned on a common cell cycle progression coordinate, where we predicted the average cell cycle dynamics of the NAD(P)H autofluorescence **(Figure 2D)** and surface area growth rate **(Figure 2E)** with Gaussian process regression. The observed oscillations match previous reports for NAD(P)H autofluorescence (Papagiannakis et al., 2017) and cell surface area growth rate (Takhaveev et al., 2023) in *S. cerevisiae*. Thus, we have confirmed the previously observed dynamics and also validated our experiment and data analysis pipeline.

**Figure 2.**
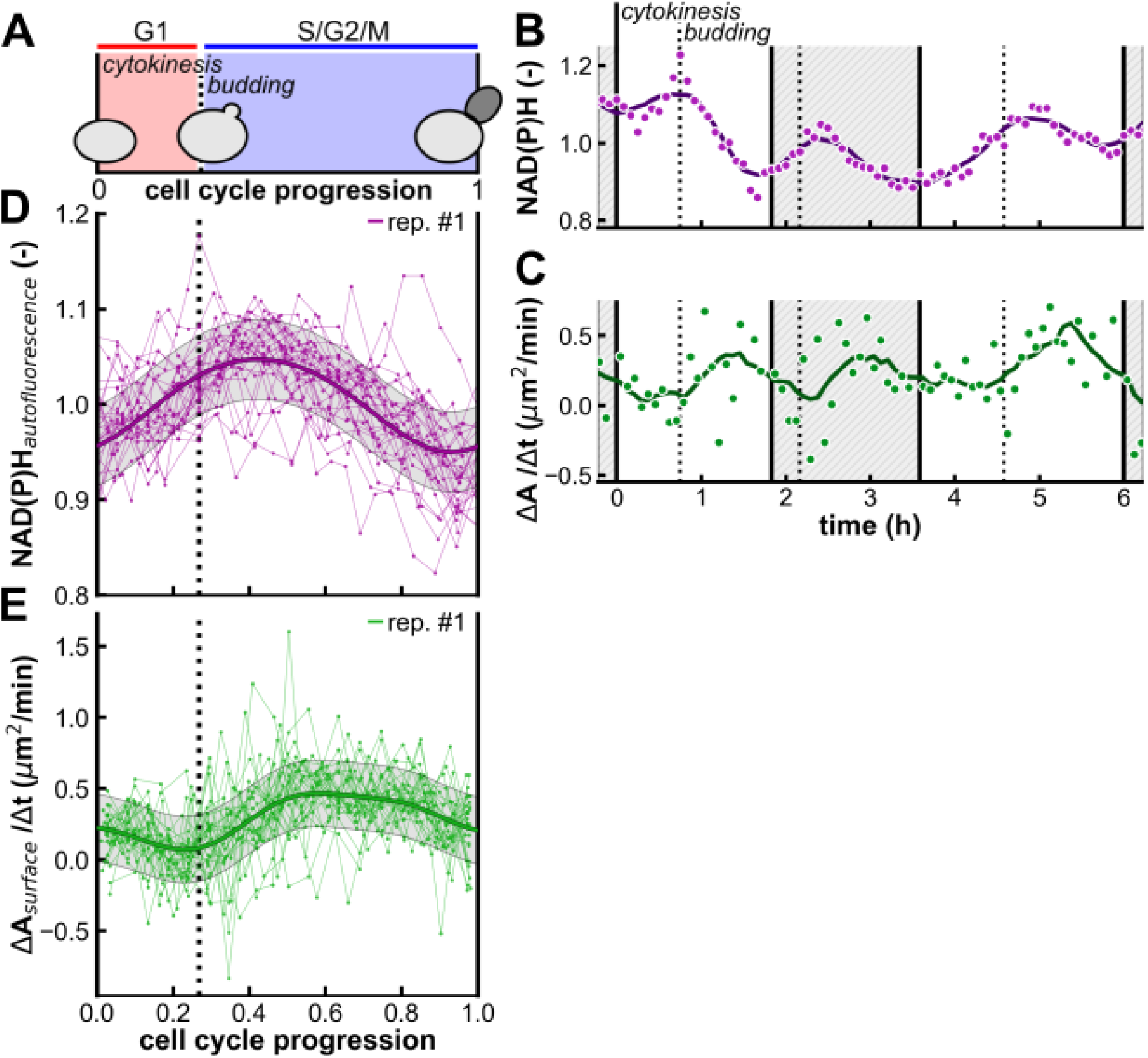
NAD(P)H autofluorescence and cell surface area growth rate oscillate over the *S. cerevisiae* cell cycle. **(A)** Single-cell trajectories were aligned for cytokinesis and budding, separating cell cycles into G1 and S/G2/M phases; **(B-C)** A single, representative cell followed over three consecutive cell cycles, showing normalised NAD(P)H autofluorescence and the cell surface area growth rate. Scatter points show raw data points and lines show smoothed trajectories. Solid and dotted vertical lines mark cytokinesis and budding, respectively. Shaded backgrounds distinguish consecutive cell cycles; **(D-E)** Average cell cycle dynamics of NAD(P)H autofluorescence and cell surface area growth rate predicted with Gaussian process regression, indicated with thick lines. A total of 32 cell cycle trajectories, shown with thin lines and scatter points, were aligned from one cytokinesis to the next, as well as for budding, which is indicated by a dotted line. Shaded area around the predicted average indicates the standard deviation.

### NAD(P)H autofluorescence and cell surface area growth rate oscillate over the *S. pombe* cell cycle

After confirming the cell cycle oscillations of NAD(P)H autofluorescence and cell surface area growth rate in *S. cerevisiae*, we asked whether similar oscillations are present in *S. pombe*. To this end, we again performed time-lapse microscopy experiments, this time using *S. pombe*, and applied our analysis pipeline. In *S. pombe*, septum ingression begins in anaphase B, but accelerates and becomes visible using bright-field imaging during telophase. At the end of telophase, the septum closes and ceases to thicken, marking septation and the M/G1 transition (Cortés et al., 2018; Pérez et al., 2016; Ramos et al., 2019). Daughter cells remain joined but septated until late S phase, when physical separation occurs (Carlson et al., 1999; Curran et al., 2022; Meister et al., 2007; Sveiczer & Horváth, 2017; Taniguchi et al., 2024; Varberg et al., 2022; Zhu et al., 2015). To align the cell cycle dynamics measured in *S. pombe* onto a common progression coordinate, we used these three events that are all identifiable in bright-field images: (i) septation, which coincides with the M/G1 transition, (ii) cell separation, occurring in late S phase and referred to as ‘cell birth’, and (iii) visible septum ingression in late anaphase. This alignment between tracked events and cell cycle phases shows that G2 covers the majority of the *S. pombe* cell cycle **(Figure 3A)**, in line with previous reports that G2 takes up approximately 70-80% of total cell cycle duration (Peng et al., 2005; Sveiczer et al., 1996).

**Figure 3.**
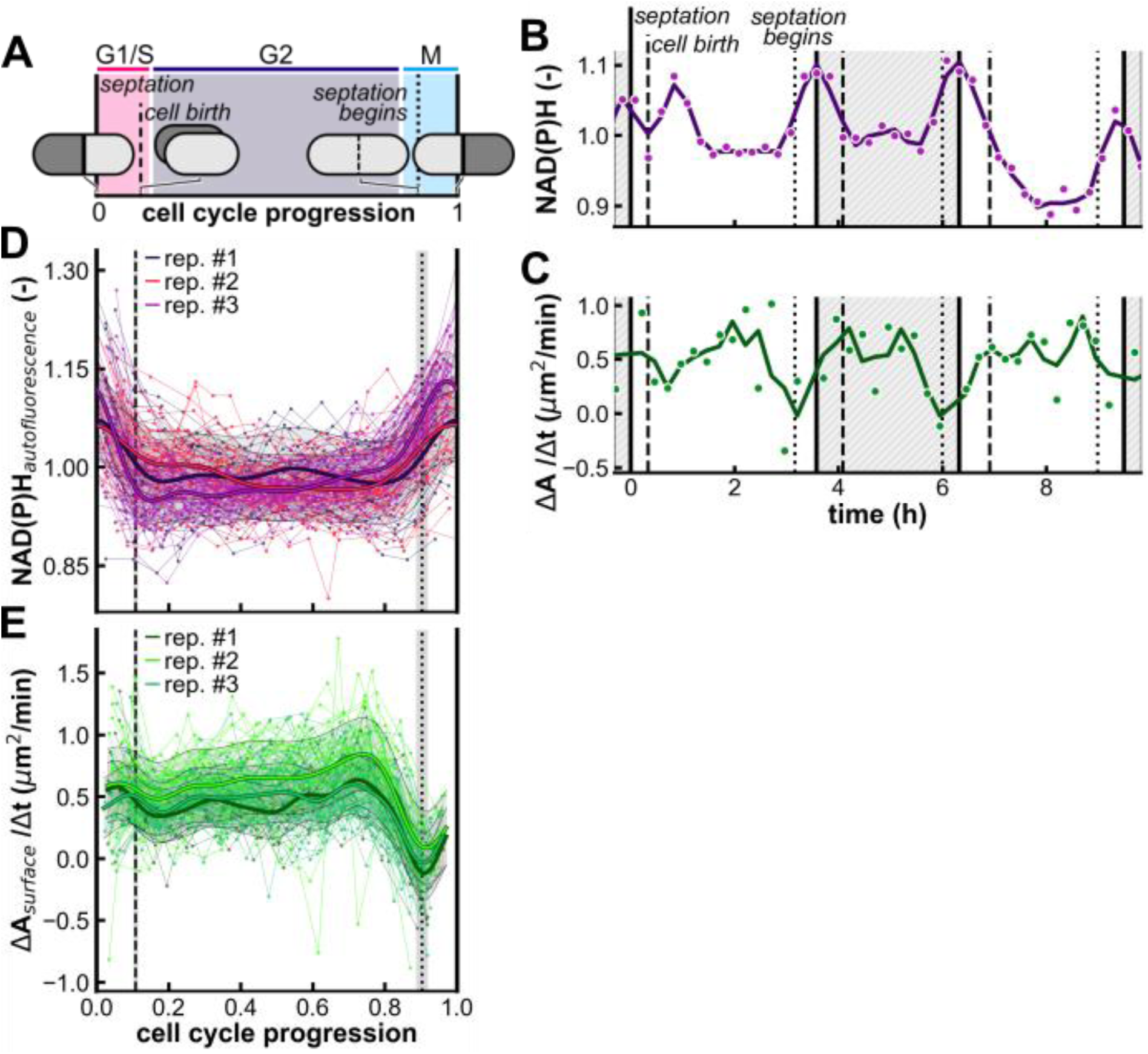
Oscillations of NAD(P)H autofluorescence and cell surface area growth rate over the *S. pombe* cell cycle. **(A)** Single-cell trajectories were aligned for septation, cell birth and initiation of septum formation, indicated with solid, dashed and dotted vertical lines, respectively. Shaded regions indicate cell cycle phases: G1/S ends shortly after cell birth, and the M phase begins shortly before the forming septum becomes visible; **(B-C)** A single, representative cell followed over three consecutive cell cycles, showing normalised NAD(P)H autofluorescence and the cell surface area growth rate. Scatter points show raw data points and lines show smoothed trajectories. Solid, dashed and dotted vertical lines indicate septation, cell birth and initiation of septation, respectively. Shaded backgrounds distinguish consecutive cell cycles; **(D-E)** Average cell cycle dynamics of NAD(P)H autofluorescence and cell surface area growth rate predicted with Gaussian process regression, shown with thick lines. 25, 46 and 55 cell cycles were analysed in the three replicate experiments. Cell cycle trajectories, shown with thin lines and scatter points, were aligned from one instance of septation to the next. Occurrence of cell birth and septum inititiation were also aligned on the cell cycle progression coordinate (dashed and dotted vertical lines, respectively). Shaded bands around the vertical lines indicate variation across three replicate experiments. Different colours denote replicates, and the shaded area around the predicted mean indicates the standard deviation.

We followed a single *S. pombe* cell over three consecutive cell cycles and observed that both the NAD(P)H autofluorescence and the cell surface area growth rate oscillate as the cell goes through the cell division cycle. NAD(P)H autofluorescence is high around septation and lower between cell birth and the initiation septum formation **(Figure 3B)**. In contrast, the cell surface area growth rate is high at the beginning of the cell cycle until it rapidly drops before septum ingression first becomes visible and stays low until septation has been completed **(Figure 3C)**. This stark decrease in growth at mitotic onset is in line with previous reports that also used time-lapse imaging of individual cells (Mitchison, 1957; Mitchison & Nurse, 1985; Sveiczer et al., 1996). Comparable oscillations to those observed in single cells were predicted with Gaussian process regression for the NAD(P)H autofluorescence **(Figure 3D)** and the cell surface area growth rate **(Figure 3E)**.

### NAD(P)H autofluorescence and cell surface area growth rate oscillate over the cell cycle in L1210 cells

Finally, we investigated whether cell cycle oscillations of NAD(P)H autofluorescence and cell surface area growth rate, as discovered in *S. cerevisiae* and *S. pombe*, also occur in murine leukaemia L1210 cells. We opted to use this cell line as L1210 cells have a short doubling time of around 10 h (Moore et al., 1966), allowing us to observe full cell cycles in experiments of a limited duration, and are spherical (Glass et al., 1980), which is convenient for automated segmentation and subsequent estimation of the cell surface area. We determined cytokinesis in the bright-field images and used it to define the beginning and end of each cell cycle **(Figure 4A)**. We also used L1210-FUCCI cells, which express fluorescenct reporters to demark the G1/S and M/G1 cell cycle transitions (Sakaue-Sawano et al., 2008; Sandler et al., 2015), in time-lapse microscopy experiments to estimate the relative duration of the G1 phase within the cell cycle. Here, we pinpointed the G1/S transition as the moment at which the degradation rate of truncated human Cdt1 tagged with mKusibara-Orange2 [mKO2-hCdt1(30/120)] was highest **(Figure S2A-B)**, since this origin licensing factor is degraded as cells enter S phase (Nishitani et al., 2001). Using this reporter strain, we determined that G1 phase covers approximately 29% of the whole cell cycle **(Figure S2C)**.

**Figure 4.**
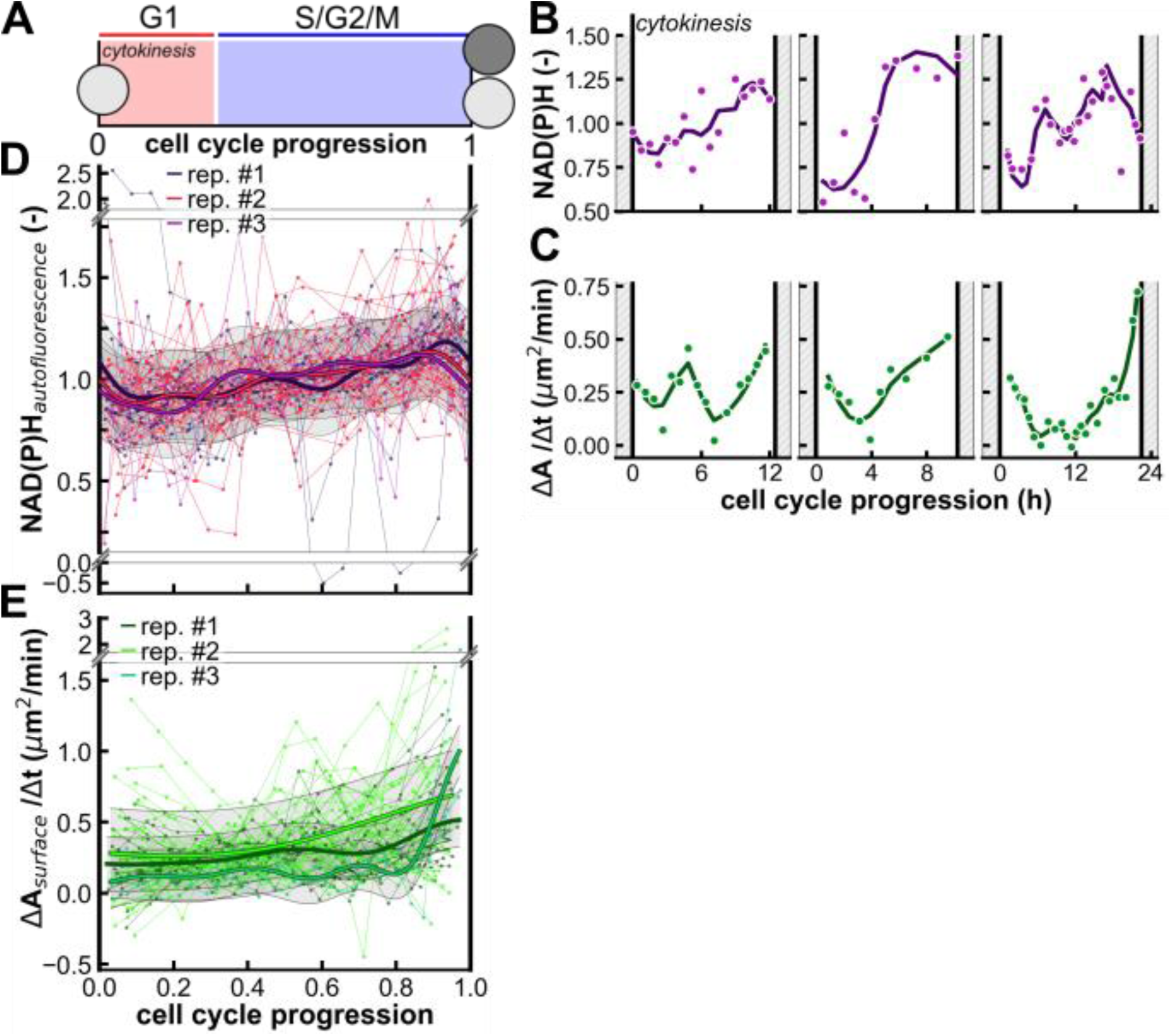
NAD(P)H autofluorescence and cell surface area growth rate oscillate over the cell cycle in L1210 cells. **(A)** Single-cell trajectories were aligned for cytokinesis, marking the beginning and end of each cell cycle; **(B-C)** Representative trajectories of normalised NAD(P)H autofluorescence and cell surface area growth rate along one cell cycle in three distinct cells. Scatter points show raw data and lines show smoothed trajectories. Vertical lines indicate cytokinesis; **(D-E)** Average cell cycle dynamics of NAD(P)H autofluorescence and the cell surface area growth rate predicted with Gaussian process regression, shown with thick lines. 27, 36 and 6 cell cycles were analysed in the three replicate experiments. Cell cycle trajectories, shown with thin lines and scatter points, were aligned from one instance of cytokinesis to the next. Different colours denote replicate experiments, and the shaded area around the predicted mean indicates the standard deviation.

First, we followed the NAD(P)H autofluorescence and the cell surface area growth rate in individual cells along one cell cycle. Despite substantial noise in the signal, we saw that the NAD(P)H autofluorescence decreases after cytokinesis to reach a minimum early in the cell cycle and then increases, until it reaches its maximum shortly before cytokinesis **(Figure 4B)**. The cell surface area growth rate slightly decreases early in the cycle and subsequently increases to reach its maximum at the end of the cell cycle **(Figure 4C)**. Next, we used Gaussian process regression to determine average cell cycle dynamics from all recorded cell cycles and found the same NAD(P)H autofluorescence dynamics that we saw on the single-cell level. There is a trough at roughly 0.11 cell cycle progression, after which the autofluorescence intensity increases until it reaches its maximum at 0.9 cell cycle progression **(Figure 4D)**. The cell surface area growth rate is low and roughly constant in the first half of the cell cycle and then increases to reach its maximum when the cell cycle ends **(Figure 4E)**. Thus, our results indicate that the cell surface area growth rate most peaks at the end of the cell cycle.

### Derivative of NAD(P)H autofluorescence and cell surface area growth rate anticorrelate in *S. cerevisiae* and *S. pombe*, but not L1210

After we determined that both the NAD(P)H autofluorescence and the cell surface area growth rate are dynamic over the cell cycle in *S. cerevisiae*, *S. pombe* and L1210 cells, we wondered whether these two signals could correlate. A link between the surface area growth rate and NADPH levels is conceivable: in *S. cerevisiae*, the cell membrane contains the majority of cellular sterols and sphingolipids and almost half of all phospholipids (Patton & Lester, 1991) and in A549 cells, the cell membrane contains 50% of all cellular lipids (Li et al., 2016). The biosynthesis of membrane building blocks, fatty acids and sterols, requires significant amounts of NADPH (Hu et al., 2017; Ratledge, 2014). Therefore, we hypothesised that faster growth of the cell surface could lead to decreasing NADPH levels, resulting in anticorrelation between the cell surface area growth rate and the derivative of the NAD(P)H autofluorescence, *i.e.*, the rate of change in NAD(P)H autofluorescence.

To test this hypothesis, we first used the Gaussian process regression predictions. Here, we calculated the average cell cycle trajectory of both NAD(P)H autofluorescence and cell surface area growth rate across replicate experiments. Subsequently, we plotted the cell surface area growth rate against the derivative of the NAD(P)H autofluorescence. We found that the cell surface area growth rate indeed globally anticorrelates with the NAD(P)H autofluorescence rate of change in *S. cerevisiae* **(Figure 5A)** and *S. pombe* **(Figure 5B)**, with Pearson correlation coefficient values of -0.94 and -0.63, respectively. In both organisms, the derivative of the NAD(P)H autofluorescence is positive when the surface area growth rate is low, suggesting that NAD(P)H levels are replenished when demand for cell membrane components is low. In contrast, the derivative of the NAD(P)H autofluorescence is negative when the cell surface area growth rate is highest, implying that the synthesis of new membrane components could result in a net decrease in NADPH levels, which would cause the measured autofluorescence to decline. However, we found no correlation between the cell surface area growth rate and the derivative of the NAD(P)H autofluorescence in L1210 cells **(Figure 5C)**. In fact, both the minimum and maximum of the cell surface area growth rate occur when the NAD(P)H autofluorescence rate of change is at comparably low values.

**Figure 5.**
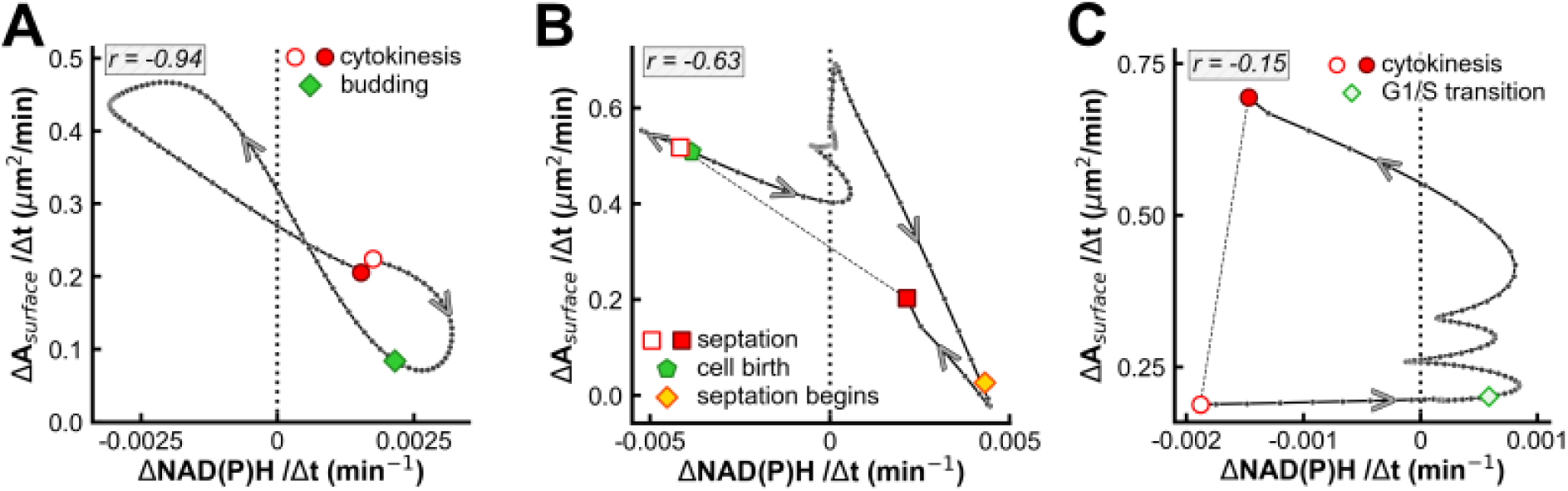
Derivative of NAD(P)H autofluorescence and cell surface area growth rate anticorrelate on the population level in *S. cerevisiae* and *S. pombe*, but not L1210. **(A-C)** Cell surface area growth rate plotted against the derivative of the NAD(P)H autofluorescence over the cell cycle in *S. cerevisae*, *S. pombe* and L1210. Cell cycle dynamics were predicted with Gaussian process regression; if replicate experiments were performed, the derivative was calculated from their averaged trajectories. Pearson correlation coefficients are shown in the top left of each panel. Black data points mark 1% cell cycle progression intervals. Coloured markers indicate cell cycle events.

After we used the Gaussian process regression outputs to demonstrate that the cell surface area growth rate and the derivative of the NAD(P)H autofluorescence anticorrelate on the population level in yeast, we next investigated if we could also detect this anticorrelation on the level of individual cell cycles. To this end, we assessed the cross-correlation between the NAD(P)H autofluorescence change rate and the cell surface area growth rate measured along individual cell cycles. Specifically, we identified the lag, expressed as a fraction of a cycle, yielding the strongest cross-correlation between the two trajectories, as well as the resulting value of the cross-correlation function. To validate the results from this analysis for all three organisms, we also generated null distributions by randomly shifting the NAD(P)H autofluorescence change rate trajectory relative to the cell surface area growth rate and compared those to the distributions obtained from the data.

We found that in *S. cerevisiae*, the lag values that yielded the strongest cross-correlation between the NAD(P)H change rate and the cell surface area growth rate centred around 0 **(Figure 6A)** and the cross-correlation peak values were negative **(Figure 6B)**, contrary to the null distributions **(Figure S3A-B)**. This result, obtained from single cell cycles, supports our finding on the population level of anticorrelation between the NAD(P)H autofluorescence change rate and the cell surface area growth rate in budding yeast. Similarly, in *S. pombe*, the lag values that yielded the strongest cross-correlation centred around 0 **(Figure 6C)** and the cross-correlation peak distribution was concentrated on negative values for the majority of cell cycles **(Figure 6D)**, contrary to the null distributions **(Figure S3C-D)**. This analysis supports an anticorrelation between the NAD(P)H autofluorescence change rate and the cell surface area growth rate in *S. pombe*. In contrast, in L1210, the lag values were spread across the full range of values **(Figure 6E)** while the cross-correlation peak values centred around 0 **(Figure 6F)**, similar to the null distributions **(Figure S3E-F)**. Consistent with our findings on the population level, these findings indicate that there is no clear correlation between the NAD(P)H autofluorescence change rate and the cell surface area growth rate in L1210.

**Figure 6.**
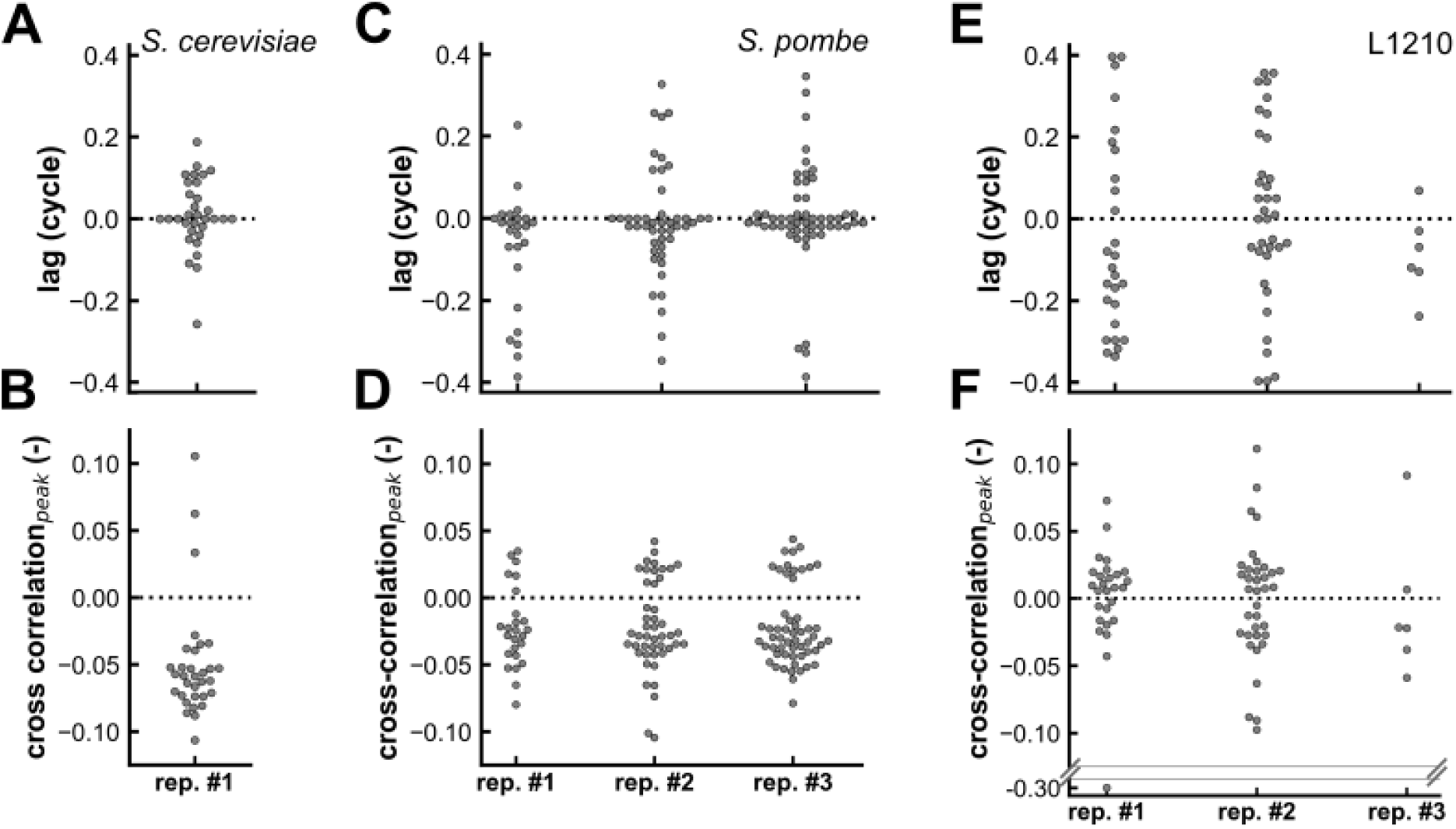
Derivative of NAD(P)H autofluorescence and cell surface area growth rate anticorrelate on the single-cell cycle level in *S. cerevisiae* and *S. pombe*, but not L1210. Cross-correlation analysis between the derivative of the NAD(P)H autofluorescence and the cell surface area growth rate in *S. cerevisiae*, *S. pombe* and L1210 reveals **(A, C, E)** the lag values yielding the strongest cross-correlation and **(B, D, F)** the resulting peak values of the cross-correlation function.

On the whole, our results show that both on the population level and on the level of individual cycles, the cell cycle dynamics of the NAD(P)H autofluorescence change rate and the cell surface area growth rate anticorrelate in budding yeast and fission yeast, but not in mammalian L1210 cells. This anticorrelation might suggest that the synthesis of membrane components, which demands large amounts of NADPH, could underly the cell cycle dynamics of the NAD(P)H autofluorescence in the two yeast species. In contrast, the synthesis of membrane components appears not to exert the same influence on the NAD(P)H levels in L1210 cells.

## Discussion

In this work, we measured two metabolism-related characteristics along the cell cycle of single cells in three organisms, without cell cycle synchronisation and without exogenous reporters. We discovered that both the cellular redox state, proxied by the NAD(P)H autofluorescence intensity, and the cell surface area growth rate, which we used as a proxy for lipid metabolism, are dynamic as cells go through the cell division cycle. Furthermore, we found that the derivative of the NAD(P)H autofluorescence anticorrelates with the cell surface area growth rate in the *S. cerevisiae* and *S. pombe*, but not L1210. Whereas the studied metabolic characteristics are dynamic in all three organisms the observed dynamics differ between organisms.

Comparing the cell cycle dynamics of the NAD(P)H autofluorescence in *S. cerevisiae*, *S. pombe* and L1210 **(Figure 7A)**, we noticed a similarity between fission yeast and the murine cells. In both, the NAD(P)H autofluorescence drops at the beginning of the cell division cycle, then increases and peaks shortly before the cell cycle ends. In contrast, the dynamics of the NAD(P)H autofluorescence in *S. cerevisiae* are almost opposite: the signal increases from the beginning of the cell cycle, reaches its maximum halfway through and subsequently decreases to reach its minimum shortly before the end of the cell cycle. The similarity between the NAD(P)H autofluorescence oscillation in *S. pombe* and L1210, which both divide symmetrically, and the striking difference to the dynamics in asymmetrically dividing *S. cerevisiae*, suggests that the type of cell division could possibly affect the dynamics of the NAD(P)H autofluorescence over the cell cycle.

**Figure 7.**
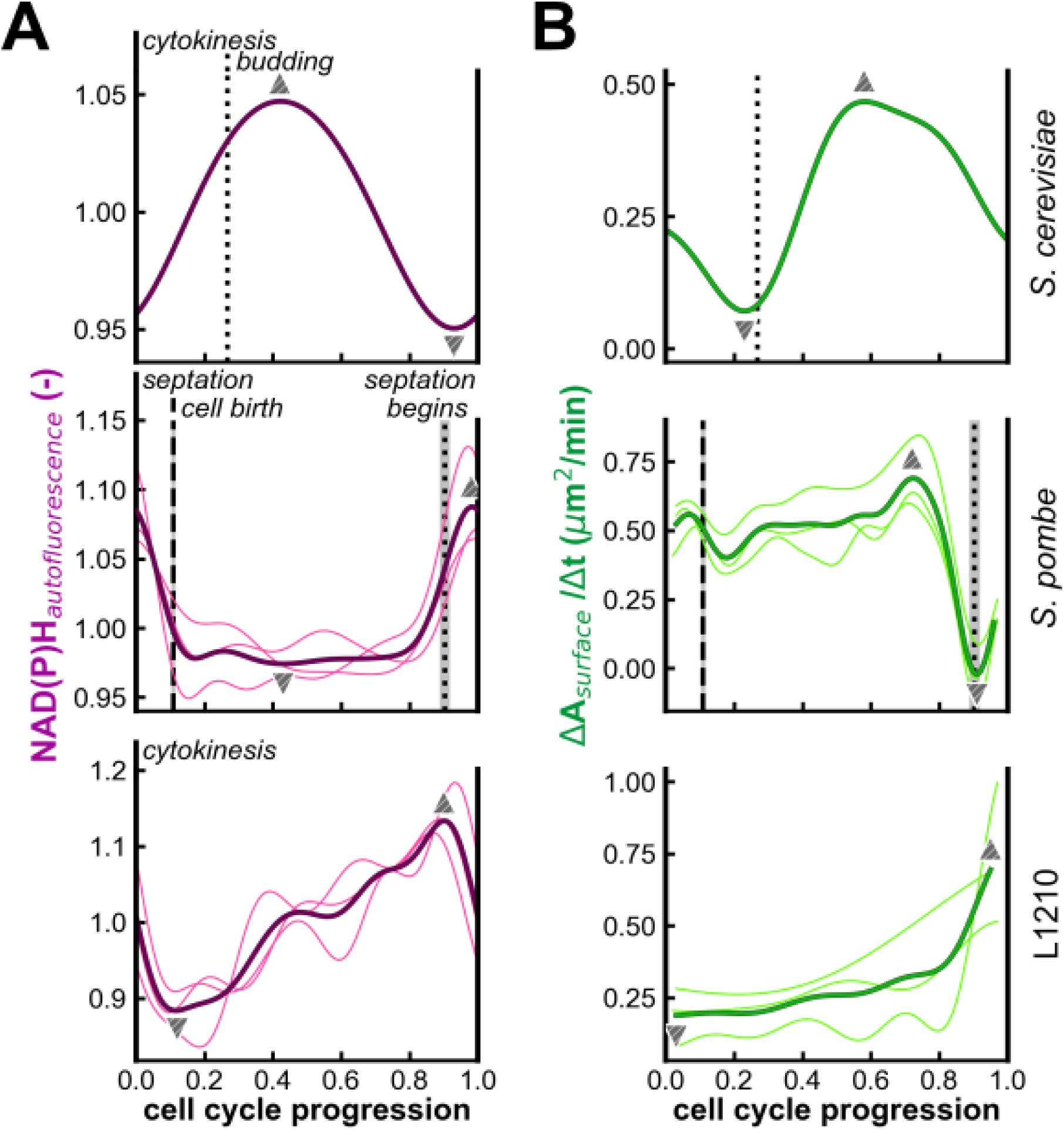
Cell cycle dynamics of NAD(P)H autofluorescence and cell surface area growth rate in *S. cerevisiae, S. pombe* and L1210. Cell cycle dynamics of **(A)** the NAD(P)H autofluorescence and **(B)** the cell surface area growth rate in the ‘average cell’, predicted with Gaussian process regression. Results for the three organisms are shown in separate rows. In each plot, thin bright lines represent replicate experiments, and thick dark lines indicate their average. Vertical lines mark cell cycle events identified from bright-field images. Individual cell cycle trajectories were aligned on a common cell cycle progression coordinate based on these events. Triangular markers denote the cell cycle progression values at which the averaged trajectories reach their minima and maxima.

Likewise, examining the cell cycle dynamics of the cell surface area growth rate in the three organisms **(Figure 7B)** there are both similarities and differences. In contrast to the similar dynamics of the NAD(P)H autofluorescence in *S. pombe* and L1210, their cell surface area growth rate dynamics are almost opposite. In L1210, the cell surface area growth rate is low at the beginning of the cell cycle and then increases from halfway through the cell cycle, peaking at the end, while in *S. pombe* the cell surface area growth rate is high for most of the cell cycle, until it drops abruptly shortly before the cell cycle ends. The dynamics in *S. cerevisiae* are yet different, decreasing during the early cell cycle with a trough around budding to subsequently increase with a peak three quarters through the cell cycle to finally decrease until the cell cycle ends. Strikingly, the decreasing cell surface area growth rate late in the cell cycle is shared between the two yeast species, which both go through closed mitosis, but not L1210, where mitosis is open, suggesting that the type of cell division could influence the cell cycle dynamics of the cell surface area growth rate.

Previous research already indicated that redox metabolism is dynamic over the cell cycle. First, the activity of the rate-limiting enzyme in the pentose phosphate pathway increases throughout the cell cycle in HeLa cells (Ma et al., 2017). This could well contribute to the increase in NAD(P)H autofluorescence we observed in L1210, as the pentose phosphate pathway is a major source of NADPH. Moreover, it has been shown that the NAD^+^/NADH ratio is cell cycle-dependent in HeLa cells. In elutriated cells, this ratio is highest in cells that are in G1, lowest in S phase cells, and has an intermediary value in G2/M phase cells (Yu et al., 2009). A more recent study on chemically synchronised cells with a notably higher time resolution has shown that the NAD^+^/NADH ratio decreases during the G1 phase, reaches a trough shortly before the G1/S transition and then increases to peak at the transition from S to G2. In contrast, the concentration of NADH increases throughout G1, peaks around the G1/S transition, and subsequently decreases to reach its minimum early in G2/M (Ahn et al., 2017). Importantly, neither the cell cycle dynamics of the NAD^+^/NADH ratio, nor those of the NADH concentration can be compared directly to the dynamics in the NAD(P)H autofluorescence we discovered. Since the excitation and emission spectra of NADPH are identical to those of NADH (Patterson et al., 2000), the autofluorescence measured in our study comes from both these reduced redox cofactors, while the two previous studies report only NAD^+^ and NADH.

Besides, previous research has also shown that the synthesis rate of lipids fluctuates along the cell cycle. In budding yeast, the cell cycle dynamics of the lipid biosynthetic activity are almost identical to those of the cell surface area growth rate (Takhaveev et al., 2023). Moreover, in chemically synchronised HeLa cells, the incorporation of multiple precursor molecules into lipids is lowest during the S phase and highest in G2/M, and the majority of acetate, *i.e.* one such precursor molecule, that is taken up during G2/M is incorporated in phospholipids (Scaglia et al., 2014). The timing of this peak in phospholipid production is in line with the peaking cell surface area growth rate we observed at the end of the L1210 cell cycle. Moreover, the peaking cell surface area growth rate shortly before cytokinesis could in part be due to mitotic swelling, a reversible increase in cell volume in mitotic cells that has been observed in L1210 (Son et al., 2015) as well as other cell lines (Zlotek-Zlotkiewicz et al., 2015). If the last recorded data point prior to cytokinesis captures a cell during mitotic swelling, its volume and surface area will be enlarged, resulting in a high cell surface area growth rate. Thus, the cell surface area growth rate dynamics reported in our study appear to be in line with previous research. However, a recent study found that the area of the plasma membrane scales approximately isometrically with cell size, and that this constant ratio is achieved by additional folding of the membrane as cells increase in size (Wu et al., 2025). For our work, we calculated the cell surface area, with the assumption that the cell is a sphere. Thus, we may underestimate the membrane area, especially as the cell cycle progresses and the cell grows and consequently, the cell surface area growth rate in L1210 might be higher than we calculated.

Overall, we have elucidated the cell cycle dynamics of the NAD(P)H autofluorescence and the cell surface area growth rate on the single-cell level in exponentially growing cells in *S. cerevisiae*, *S. pombe* and L1210. Our results show that both redox and lipid metabolism are dynamic along the cell cycle in all three organisms, which suggests that metabolic oscillations along the cell cycle could well be a conserved characteristic among eukaryotes. This is in line with previous research that has highlighted the importance of redox balance (H.-J. Kang et al., 2018; Kim et al., 2011) and lipid metabolism (Al-Feel et al., 2003; Scaglia et al., 2014; Takemoto et al., 2016) for cell cycle progression during mitosis and linked lipid synthesis during mitosis to nuclear envelope expansion (Khondker et al., 2024; Makarova et al., 2016; Rodriguez Sawicki et al., 2019). Further research building on the current results should investigate how these metabolic oscillations are orchestrated and how they impact or even drive cell cycle progression.

## Materials and methods

### Construction of PDMS wells

To perform time-lapse imaging of L1210 cells, they were cultured in PDMS wells crafted from a 5 mm thick slab of PDMS that was prepared as described previously (Huberts et al., 2013) in a glass petri dish (ø 12 cm). From this slab, each individual well was created by first cutting a rectangle of approximately 1.7 cm by 2 cm and then cutting and removing a square of 0.9 cm by 1 cm from its middle. The resulting well was attached to a glass cover slip after activation by UV treatment as reported (Huberts et al., 2013).

### Cell lines and culturing

Stocks of *Saccharomyces cerevisiae* YSBN6, derived from the S228C background (Canelas et al., 2010) and *Schizosaccharomyces pombe* CBS356 were obtained from the lab inventory. L1210 wild-type and L1210-FUCCI (Sandler et al., 2015) cell lines were gifted by the Itamar Simon lab.

*S. cerevisiae* and *S. pombe* were cultured in Verduyn minimal medium (Verduyn et al., 1992) buffered at pH 5.0 with 10 mM potassium phthalate and with 2% glucose as a carbon source. To grow *S. pombe*, the medium was further supplemented with 1 mg/mL citrate from tri-sodium citrate dihydrate, *i.e.* 1.56 mg/mL Na_3_C_6_O_7_H_5_ ·2 H_2_O. Cells were pre-cultured in 10 mL of medium in 100 mL flasks at 30°C under constant rotation at 300 rpm. Prior to time-lapse microscopy experiments, cultures were maintained in the exponential phase for at least 12 h prior to the experiment through repeated dilution. Before the experiment, cultures were diluted to OD_600_ 0.1 and were grown for 2 h more, after which 1-4 µL of cells were pipetted onto a glass cover slip and a 1% agarose pad containing the growth medium was placed on top. To limit drying of the agar pad over the course of the time-lapse experiment, a drop of 100 µL sterile medium was pipetted on the glass next to the agar pad and a second cover slip was placed on top of the cover slip holder to create a closed environment and limit evaporation.

L1210 wild-type and FUCCI cells were grown in RPMI 1640 medium (Gibco) with 10% fetal bovine serum (FBS; Gibco), 11 mM D-glucose, 2 mM L-glutamine and 0.01 mM phenol in 75 cm^2^ culture flasks with filter screw caps (TPP®) at 37 °C in an incubator with humidified atmosphere of 5% CO_2_ and 95% air. The medium was changed at least every 48 hours and cells were routinely seeded to a density of 0.5·10^5^ cells/mL, based on cell counts obtained with a Fast-Read 102 haemocytometer (Biosigma) after staining with 0.4% Trypan Blue Solution. For time-lapse microscopy, cells were grown to a density of 1-2·10^6^ cells/mL and were then spun down for 6 min at 900 g. Subsequently, they were resuspended to yield a density of 1·10^5^ cells/mL, using RPMI medium with 10% FBS, 11 mM D-glucose, 2 mM L-glutamine and 0.01 mM phenol red and buffered with 25 mM HEPES to counteract acidification of the medium over the course of the time-lapse experiment. The resuspended cells were incubated at 37 °C in a humidified atmosphere of 5% CO_2_ and 95% air for 6 h. Subsequently, a PDMS well was completely filled up with approximately 0.5 mL of wild-type L1210 culture and covered with a glass cover slip to limit medium evaporation during the time-lapse experiment. Alternatively, L1210-FUCCI cells were loaded into glass bottom µ-Slide 8 well chambers (ibidi), which were sealed with parafilm to minimise evaporation of the culture medium.

### Image acquisition

Time-lapse microscopy images of *S. cerevisiae*, *S. pombe* and wild-type L1210 were obtained on a Nikon Eclipse Ti-E inverted wide-field fluorescence microscope fitted with the Nikon Perfect Focus System (PFS) to maintain focus, an Andor-DU-897 EX camera with the readout mode set to 1 MHz without gain amplification and Nikon S Fluor Oil Iris objectives (NA = 1.3) with ×100 magnification for *S. cerevisiae* and *S. pombe* and ×40 magnification for L1210. An incubator box (Life Imaging Services) kept the setup at a constant temperature of 30 °C for *S. cerevisiae* and *S. pombe* or 37 °C for L1210 cells. For bright-field images, light came from a halogen lamp fitted with a 420 nm beam-splitter filtering out all shorter wavelengths. To image NAD(P)H autofluorescence, excitation light at 360 nm was provided by a Lumencor AURA excitation system and a 350/50 nm band-pass filter, 409 nm beam splitter and 435/40 nm emission filter were used. For *S. cerevisiae* and L1210, excitation light intensity was set to 3% with an excitation time of 200 ms. For *S. pombe*, to achieve a sufficiently high signal-to-noise-ratio, stronger excitation was used at 6% with an exposure time of 400 ms.

Time-lapse microscopy data of L1210-FUCCI cells were acquired on a similar microscope setup, but equipped with a 2xAndor LucaR EM-CCD camera and a CoolLED pE-2 excitation system. Geminin-mAG was imaged in the YFP channel, with excitation at 500 nm, with a 520/20 nm band-pass filter and a 515 nm beam splitter as well as a 535/30 nm filter for the emitted light. The excitation light intensity was set at 9% and the exposure time was 200 ms. mKO2-hCdt1(30/120) was imaged in the RFP channel, with excitation at 565 nm, a 560/40 nm band-pass filter, a 585 nm beam splitter and a 630/75 nm emission filter. Light intensity was set at 9% and an excitation time of 50 ms was used.

For *S. cerevisiae*, both bright-field and NAD(P)H autofluorescence images were acquired every 5 min. For *S. pombe*, the imaging interval was 5 min for bright-field and 15 min for NAD(P)H autofluorescence, which was thus imaged at every third time point recorded in the bright-field channel. This approach limited the phototoxicity associated with the higher dose of excitation light compared to *S. cerevisiae*. Wild-type L1210 cells were imaged every 15 min in the bright-field and every third frame was also imaged in the NAD(P)H autofluorescence channel, resulting in a 45 min imaging interval. Moreover, wild-type L1210 cells were first imaged only in the bright-field channel for the first 16 hours to make sure cells were dividing as expected and there was no bacterial contamination in the culture. After this initial time-lapse, the cells were imaged for an additional 30-40 hours in both the bright-field and the NAD(P)H autofluorescence channel. For L1210-FUCCI cells, the imaging interval also was 15 minutes, but fluorescence images were acquired at every time point and from the beginning of the time-lapse.

### Cell segmentation

To obtain cell masks, only the bright-field images for which a corresponding NAD(P)H autofluorescence image was available were segmented with YeaZ (Dietler et al., 2020) using its standard settings. All obtained masks were inspected and manually corrected if necessary. Especially for *S. pombe*, images obtained around septation often required manual correction. Before septation had been completed, YeaZ occasionally segmented the mother cell as two separate daughters. Conversely, after septation, YeaZ still segmented some daughter cells together, as if they were one.

### Calculating cell surface area and determining surface area growth rate

YeaZ fits an ellipse to every cell mask it creates and provides the lengths of its major and minor axes (D and d, respectively). These axis lengths were converted into the corresponding major radius (R) and minor radius (r), which in turn were used to calculate the cell surface area of every cell. For *S. cerevisiae*, the cell was assumed to be a prolate ellipsoid with its surface area described by

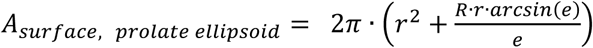

in which

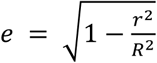

For *S. pombe* a rod shape was assumed and its surface area was calculated with

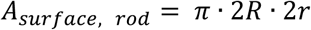

L1210 cells were assumed to be spherical with a surface area that was calculated based on the average of the major and minor radius using

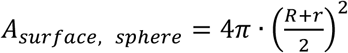

The surface area growth rate was calculated by determining the difference in cell surface area between two consecutive data points and dividing this value by the time that elapsed between the two measurements. Notably, before calculating the surface area growth rate of L1210 cells, to counteract noise stemming from small imperfections in the cell masks, the cell surface area trajectories were smoothed with a Savitzky-Golay filter, fitting a second-order polynomial and using a window size of 7.

### Background correction

To correct for background fluorescence stemming from the medium in the obtained NAD(P)H autofluorescence images, the modal pixel intensity value of each image was subtracted from each individual pixel intensity within that frame. However, for two L1210 time-lapse experiments with higher confluency, an alternative approach had to be used. Due to the high cell density in these experiments, the modal pixel intensity value corresponds to the pixel intensity most common inside cells instead of the background intensity outside cells. Therefore, for these two replicate experiments, instead of the modal pixel intensity value, the minimum pixel intensity was used for background correction. First, the average of the minimum pixel intensity values across all positions was calculated for every time point. The resulting series of values was smoothed with the Savitzky-Golay filter, fitting a second-order polynomial and using a window size of 17. This approach yielded a background intensity value for every time point. Subsequently, the relevant background intensity value was subtracted from all pixel intensity values to background correct every image. However, subtracting the minimum instead of the modal pixel intensity value caused an overestimation of the average fluorescence intensity inside the cell. If not corrected, we would underestimate the relative changes in fluorescence over the course of a cell cycle. To compensate this difference between the minimum and the modal pixel intensity value, we also subtracted a constant correction value from every average cellular fluorescence intensity value. This constant was chosen such that, after correction, the average cellular fluorescence intensity measured across the whole experiment was equal to 12 AU, which was the average fluorescence intensity in cells measured after background correction in the replicate corrected by subtraction of the modal pixel intensity value.

### Data detrending

To remove an increase in average NAD(P)H autofluorescence on the multi-cell cycle time scale that is likely associated with aging, *S. cerevisiae* and *S. pombe* data were detrended. Specifically, linear regression was performed, yielding a function that describes average cellular fluorescence intensity as a function of time. NAD(P)H autofluorescence data were detrended by dividing the average cellular fluorescence by the time-dependent fluorescence intensity value predicted with linear regression. Notably, data from L1210 were not detrended in this manner, since cells were typically only followed for a single cell cycle early in the time-lapse experiment.

### Cell cycle alignment

Cell cycle events visible in the bright-field images were detected and tracked by hand. Cytokinesis and budding were detected in *S. cerevisiae*. In *S. pombe*, cell birth, *i.e.* the moment the two daughter cells separate from each other, the initial appearance of the forming septum, and septation were tracked. In L1210 cells, only cytokinesis was detected.

To combine data from all individually observed cell cycles, they were aligned for cell cycle progression on a common time coordinate. Here, 0 and 1 represent the beginning and end of the cell cycle, coinciding with cytokinesis in *S. cerevisiae* and L1210, and septation in *S. pombe*. The occurrence of other cell cycle events on this relative time coordinate, *e.g.* budding in *S. cerevisiae*, was determined as follows. For every cell cycle, the time elapsed between the beginning of the cycle and the occurrence of the event was divided by the total duration of that cycle. Once all cell cycles had been assessed, the average cell cycle progression value corresponding to occurrence of the event was calculated. Subsequently, we determined the timing of all other data points, that did not coincide with the occurrence of a cell cycle event. We first split each cycle into subphases starting and ending with a tracked cell cycle event. Next, we placed the data points that coincided with a tracked event at the determined cell cycle progression values. Finally, we obtained the cell cycle progression values for all data points between the two events by evenly dispersing them over the interval bounded by those cell cycle events. Repeating this approach for all cell cycles, we assigned cell cycle progression values to all data points.

### Gaussian process regression

The aligned single-cell data were used to predict the average cell cycle dynamics of the NAD(P)H autofluorescence and the cell surface area growth rate with Gaussian process regression. Specifically, we used a radial basis function (RBF) kernel as a prior and a white kernel with free noise level, and maximised the log-marginal likelihood.

Before applying Gaussian process regression, we normalised each NAD(P)H autofluorescence cell cycle trajectory to its own mean in order to limit the influence of cell-to-cell variation in fluorescence intensity on our analysis. Furthermore, we ensured that the mean NAD(P)H signal at the beginning and end of the cell cycle in the Gaussian process output were identical to each other. If Gaussian process regression was applied to data corresponding to an isolated cell cycle, the transition would not be assessed. To solve this issue, we simulated three consecutive cell cycles by duplicating our cell cycle-aligned data and transforming the cycle progression values by subtracting or adding 1, to simulate the previous and the next cell cycle, respectively. Then, Gaussian process regression was applied, predicting the average signal of the NAD(P)H autofluorescence along three consecutive cycles. The progression of the signal across the cell cycle transition was assessed explicitly for the middle cycle. Therefore, we selected the Gaussian process regression results corresponding to the middle cycle, with matching NAD(P)H autofluorescence signal at the beginning and end.

### Cross-correlation analysis

The aligned single-cell data were used in a cross-correlation analysis to assess the relation between the derivative of the NAD(P)H autofluorescence and the cell surface area growth rate in individual cell cycles. First, we interpolated the data such that there were 99 data points between cell cycle progression values 0 and 1, resulting in a timestep of 0.01 cycle between data points. We then subtracted the mean of the trajectory from every data point, so that both interpolated trajectories were centred around 0. Next, we calculated the cross-correlation between the normalised interpolated trajectories. We assessed the cross-correlation function across a range of lag values from -0.4 cycle to 0.4 cycle and identified the lag value that yielded the largest absolute value of the cross-correlation function, *i.e.* the strongest cross-correlation. Hereby, we obtained the peak value of cross-correlation between the NAD(P)H autofluorescence change rate and the cell surface area growth rate as well as the lag value.

As a control, we generated null distributions of the lag and the cross-correlation peak value. Here, we shifted the interpolated derivative of the NAD(P)H autofluorescence by a randomly selected shift. Potential shift values ranged from -0.4 cycle to 0.5 cycle, with a step size of 0.1 cycle between potential shifts and excluded 0. We then used the shifted trajectory of the NAD(P)H autofluorescence change rate to quantify cross-correlation with the unshifted cell surface area growth rate trajectory using the approach described above.

## Acknowledgements

The wild-type L1210 and L1210-FUCCI cell lines were a kind gift of the Itamar Simon lab. We thank Andreas Milias-Argeitis for his help with the cross-correlation analysis.

## SUPPLEMENTAL MATERIAL

**Figure S1.**
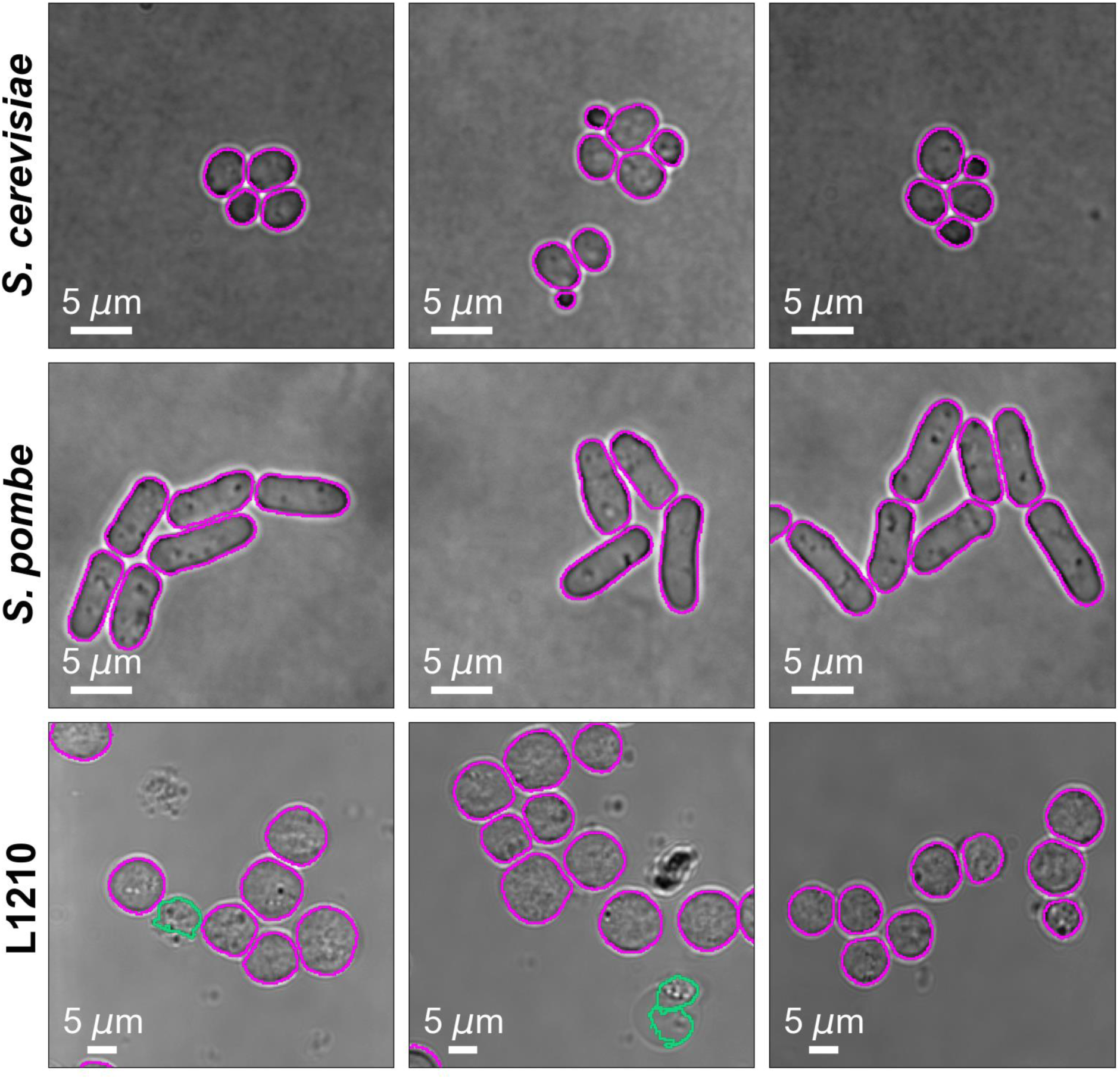
YeaZ faithfully segments bright-field images of *S. cerevisiae, S. pombe* and L1210. YeaZ (Dietler et al., 2020) was used to segment bright-field images of the yeasts *S. cerevisiae* and *S. pombe* (100× magnification) and murine leukaemia L1210 cells (40× magnification). For each organism, three representative images are shown. Outlines of the cell masks obtained with YeaZ are shown in bright pink. For L1210, some dead, blebbed cells were segmented as well; outlines of these cell masks are indicated in green.

**Figure S2.**
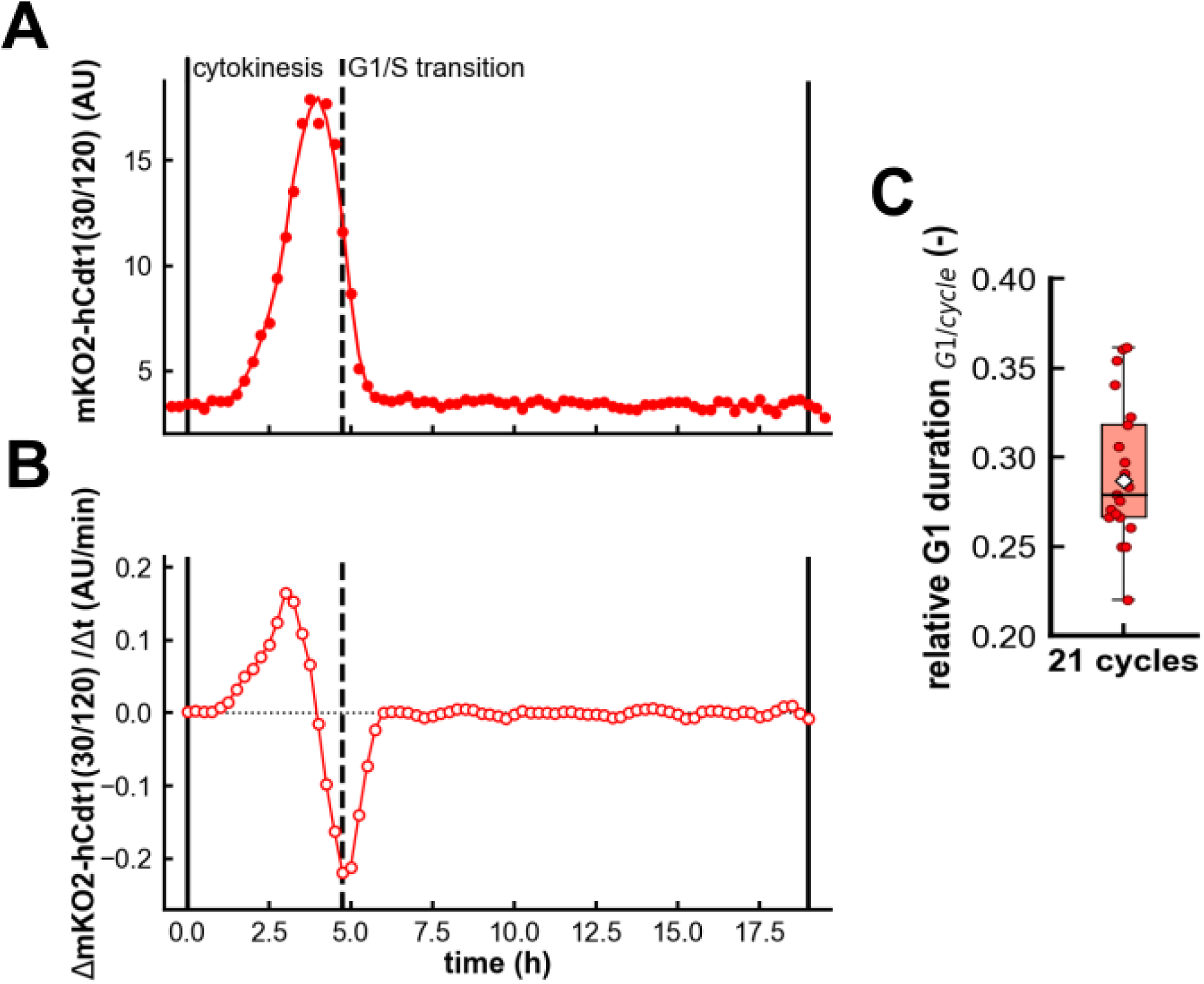
Determination of G1 phase duration in L1210-FUCCI cells. **(A)** Average fluorescence intensity of truncated human Cdt1 tagged with mKusibara-Orange2 [mKO2-hCdt1(30/120)] along the cell cycle in a representative L1210-FUCCI cell. Scattered data points represent the raw data; the line shows the smoothed trajectory. The moment of cytokinesis was detected in bright-field images. mKO2-hCdt1(30/120) fluorescence intensity increased over the G1 phase and dropped abruptly when the cell entered S phase; **(B)** Derivative of the smoothed trace in A, which was used to detect the G1/S transition. Since the origin licensing factor Cdt1 is degraded when cells enter S phase, we determined when the derivative of the mKO2-hCdt1(30/120) signal was minimal and thereby detected the G1/S transition; **(C)** Relative duration of the G1 phase in 21 L1210-FUCCI cells; the white diamond represents the mean value. The relative G1 duration, *i.e.* the fraction of the cell cycle that is taken up by the G1 phase, was determined by dividing the duration of the G1 phase by the duration of the whole cell cycle.

**Figure S3.**
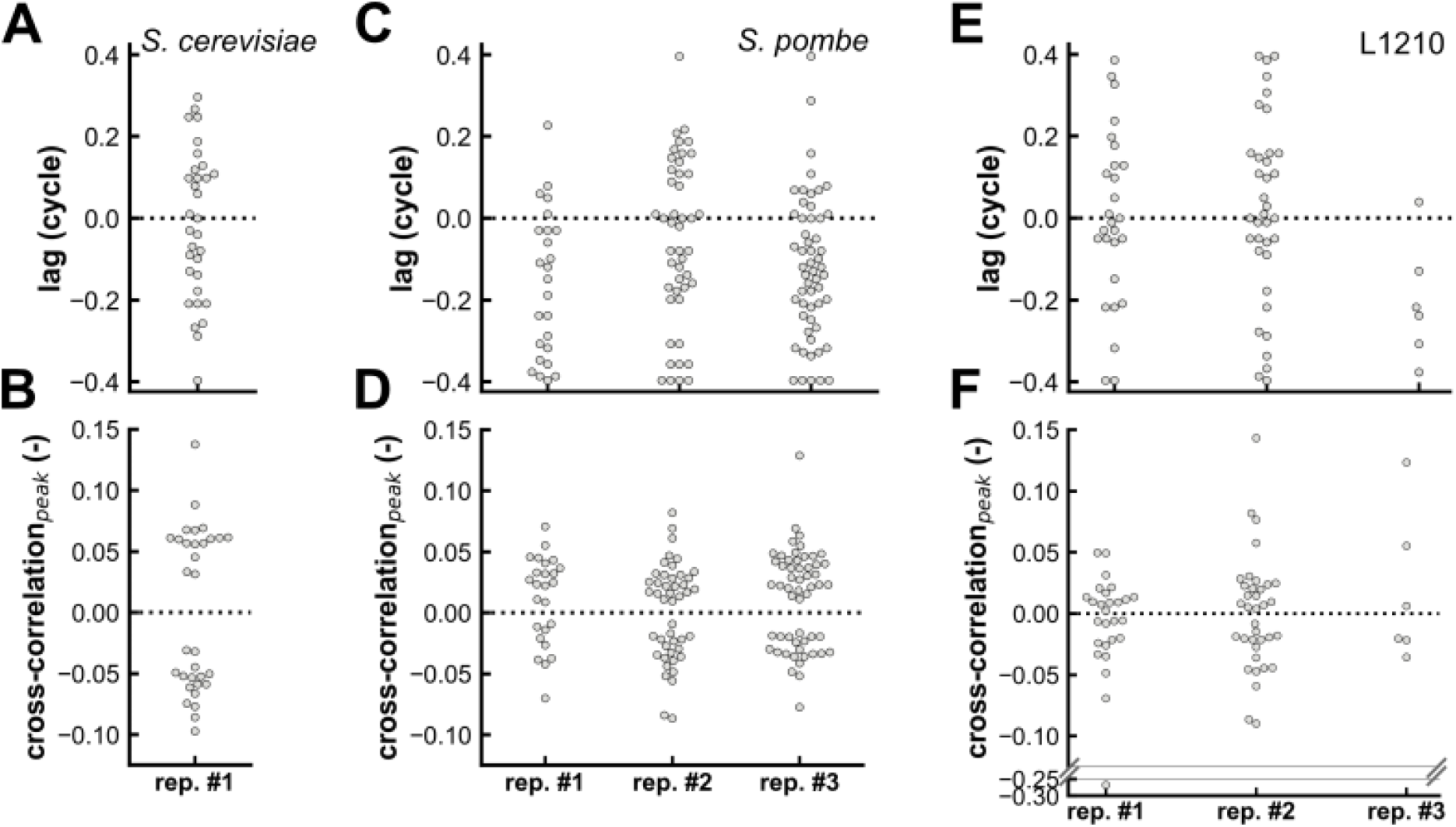
Null distributions of lag values and peak values for cross-correlation between the NAD(P)H autofluorescence change rate and the cell surface area growth rate in *S. cerevisiae, S. pombe* and L1210. The single-cell cycle trajectory of the NAD(P)H autofluorescence was randomly shifted after which cross-correlation analysis was performed to find **(A, C, E)** the lag value that yielding the strongest cross-correlation with the cell surface area growth rate and **(B, D, F)** the corresponding peak values of the cross-correlation function.

